# Inactivation of face selective neurons alters eye movements when free viewing faces

**DOI:** 10.1101/2023.06.20.544678

**Authors:** Reza Azadi, Emily Lopez, Jessica Taubert, Amanda Patterson, Arash Afraz

## Abstract

During free viewing, faces attract gaze and induce specific fixation patterns corresponding to the facial features. This suggests that neurons encoding the facial features are in the causal chain that steers the eyes. However, there is no physiological evidence to support a mechanistic link between face encoding neurons in high-level visual areas and the oculomotor system. In this study, we targeted the middle face patches of inferior temporal (IT) cortex in two macaque monkeys using an fMRI localizer. We then utilized muscimol microinjection to unilaterally suppress IT neural activity inside and outside the face patches and recorded eye movements while the animals free viewing natural scenes. Inactivation of the face selective neurons altered the pattern of eye movements on faces: the monkeys found faces in the scene but neglected the eye contralateral to the inactivation hemisphere. These findings reveal the causal contribution of the high-level visual cortex in eye movements.

**Significance:** It has been shown, for more than half a century, that eye movements follow distinctive patterns when free viewing faces. This suggests causal involvement of the face-encoding visual neurons in the eye movements. However, the literature is scant of evidence for this possibility and has focused mostly on the link between low-level image saliency and eye movements. Here, for the first time, we bring causal evidence showing how face-selective neurons in inferior temporal cortex inform and steer eye movements when free viewing faces.

## Introduction

In the retina, high concentration of photoreceptors can be found only in the fovea, thus the most efficient way for the visual system to gather information is to move the eyes to survey the visual field. Unguided scanning of the visual field is inefficient, thus the oculomotor system needs to rely on the incoming visual information to steer the eyes. As a result, a bidirectional causal motif forms: the input to the eyes guides the eye movements, which in turn select the input to the eyes.

How does the visual input direct eye movements? Saliency maps have been proposed to predict the location of future fixations by weighting the features in a scene according to their visual saliency (1). Earlier models of saliency maps relied solely on low-level visual properties such as brightness, contrast, and spatial frequency (2–4). However, high-level visual properties such as complex object features also affect eye movement behavior. Therefore, recent saliency map models have begun to incorporate complex features to improve their predictive power (5–9). These models suggest functional connectivity between parts of the brain that process high-level object information and the oculomotor system. Nevertheless, there is currently no neurophysiological evidence supporting this idea.

Complex object features are processed in high-level visual areas such as inferior temporal (IT) cortex (10–12). Therefore, we hypothesized that neural activity in IT cortex should play a causal role in controlling eye movements. To investigate this hypothesis, we chose to study the link between eye movements when free viewing faces and face-selective neurons in IT cortex (13, 14). There are a number of reasons for this choice. First, determining the preferred stimulus of a group of cells in IT cortex is less speculative for the case of face-selective neurons. Second, these neurons are typically clustered, making it possible to perturb the activity of a group of cells with similar function (13, 15). Third, faces have geometrical regularities that facilitate quantitative analysis of the results (16). Last but not least, the natural pattern of eye movements when viewing faces are relatively standard and distinct from other objects, thus can be easily quantified and tracked.

The specific patterns of eye movements during free viewing of faces was first noticed by Yarbus (17) and consistently reported since (18–21). In ‘eye movement and vision’ (17) Yarbus describes: “…the faces of the people shown in the picture attract the observer’s attention much more than the figures…” and then later “…when looking at a human face, an observer usually pays most attention to the eyes, the lips, and the nose.” This pattern of eye movement has also been reported in nonhuman primates (22), and we will refer to it as Yarbus T (23).

Low-level saliency models may partially explain this behavior because eyes and mouth typically have higher visual contrast (24), nevertheless, possible involvement of high-level visual processing in steering the eye movements while viewing faces and complex objects cannot be rejected.

Here we aimed at testing the role of IT face-selective neurons in eye movements by causal manipulation of their function. We used muscimol, a strong GABA_A_ agonist (25) and potent neural silencer, to reversibly inactivate clusters of face-selective neurons, as well as control regions unilaterally in IT cortex of two macaque monkeys (Figure 1). We then investigated the resulting effects on eye movements when free viewing faces and other objects. We chose to target the middle face patch in the face processing network because it contains the largest number of face selective neurons, it can be reliably localized and it is hypothesized to be homologous to the human fusiform face area (FFA) (13, 26–30). Face selective cells retain some retinotopy and are shown to causally contribute to face recognition behavior only in their contralateral visual field (31), thus we chose to target only one hemisphere at a time. Using this design, we can compare the behavioral results across the two hemifields within each injection session, a feature that increases the statistical power of the study. Moreover, given the logistic limitations of performing microinjection in a deep brain structure like IT cortex (see Methods), unilateral injections are much safer for the animal.

**Figure 1:**
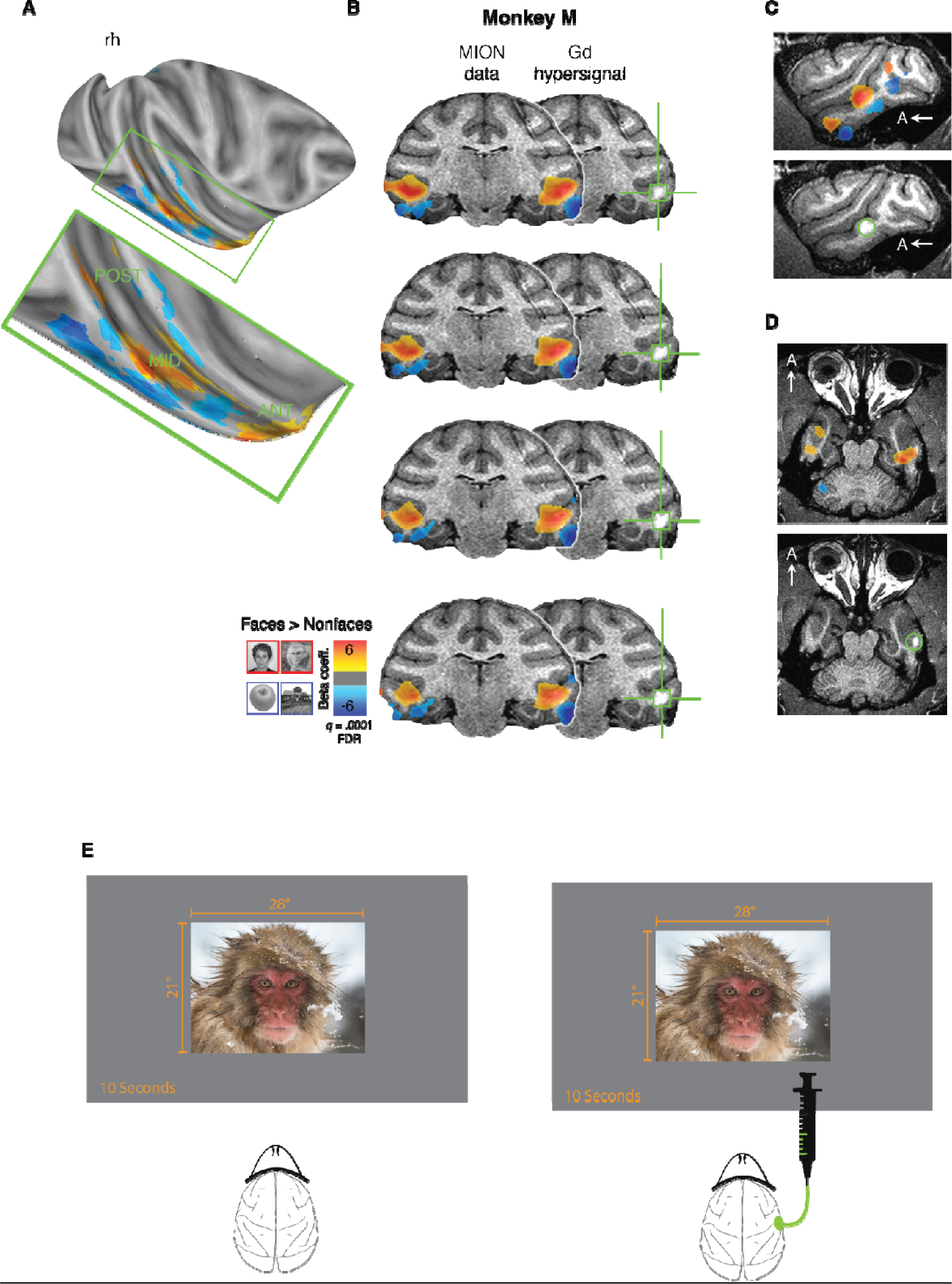
Study overview. (A) Functional localizer data from Monkey Mr projected onto a partially inflated cortical surface, showing a lateral view of the right hemisphere. The voxelwise statistical threshold was set at q = 0.001 (corrected for False Discovery Rate). The localization procedure successfully identified multiple face-selective patches, primarily in the posterior (POST), middle (MID), and anterior (ANT) regions of the superior temporal sulcus. (B-D) Post-Gadolinium-DTPA infusion anatomical scans with Beta maps superimposed, illustrating the contrast between face stimuli (hot colors) and nonface stimuli (cool colors) to validate the targeting procedure. The bright white hypersignal is associated with the infusion. (B) Coronal slices with 1 mm separation, starting approximately +2 mm anterior to the interaural line and presented posterior to anterior (top to bottom). The green crosshairs indicate the signal from injected Gadolinium-DTPA. (C-D) Sagittal and transverse views. A = anterior. (E) Monkeys’ eye movement was recorded during free-viewing of visual stimuli presented on a screen subtending 28° x 21° during baseline sessions and after injection of 5 or 10 µL in the middle face patch of the left or the right hemisphere.

## Results

In this experiment, two male macaque monkeys (*macaca mulatta*) were used. Each trial began with the animals fixating on a central fixation point for 1 second before an image (28 x 21 dva) was displayed on the screen. The image was chosen randomly from a set of 62 images, consisting of human and nonhuman primate faces, objects, plants and scenes. The image was displayed for 10 seconds on the screen. During the image presentation the animals were not required to hold fixation, instead they could freely move their eyes and were rewarded for keeping their gaze within the image boundaries every 1 to 3.5 seconds. We used an fMRI face/object localizer to locate the middle face patch in IT cortex (13, 32, 33) (Figure 1A-D), as well as control areas with no face selectivity, for muscimol microinjections. These microinjections were performed through a cranial chamber. We also recorded the animals’ natural behavior without any injection in multiple separate baseline sessions interleaved with injection sessions. We employed two different volumes of muscimol: 5 µL and 10 µL, which were injected into the middle face patch and control regions in separate injection sessions.

### Natural viewing patterns for faces

First, we aimed at systematic documentation of the monkeys’ natural free viewing patterns for the faces in our study. Figure 2 provides examples of average fixation heatmaps on both monkey and human faces across all baseline trials. During these trials, the animals consistently identified and fixated on faces, doing so in 98.9% and 98.7% of the trials for monkeys Mr and Fd, respectively, with a reaction time of only 695 ms and 418 ms. Additionally, our results indicated that the majority of the looking time in each trial was on faces, accounting for 68.1% and 72.1% of the total trial time for monkeys Mr and Fd, respectively. This establishes that similar to human observers, monkeys prefer to spend more time looking at faces.

**Figure 2:**
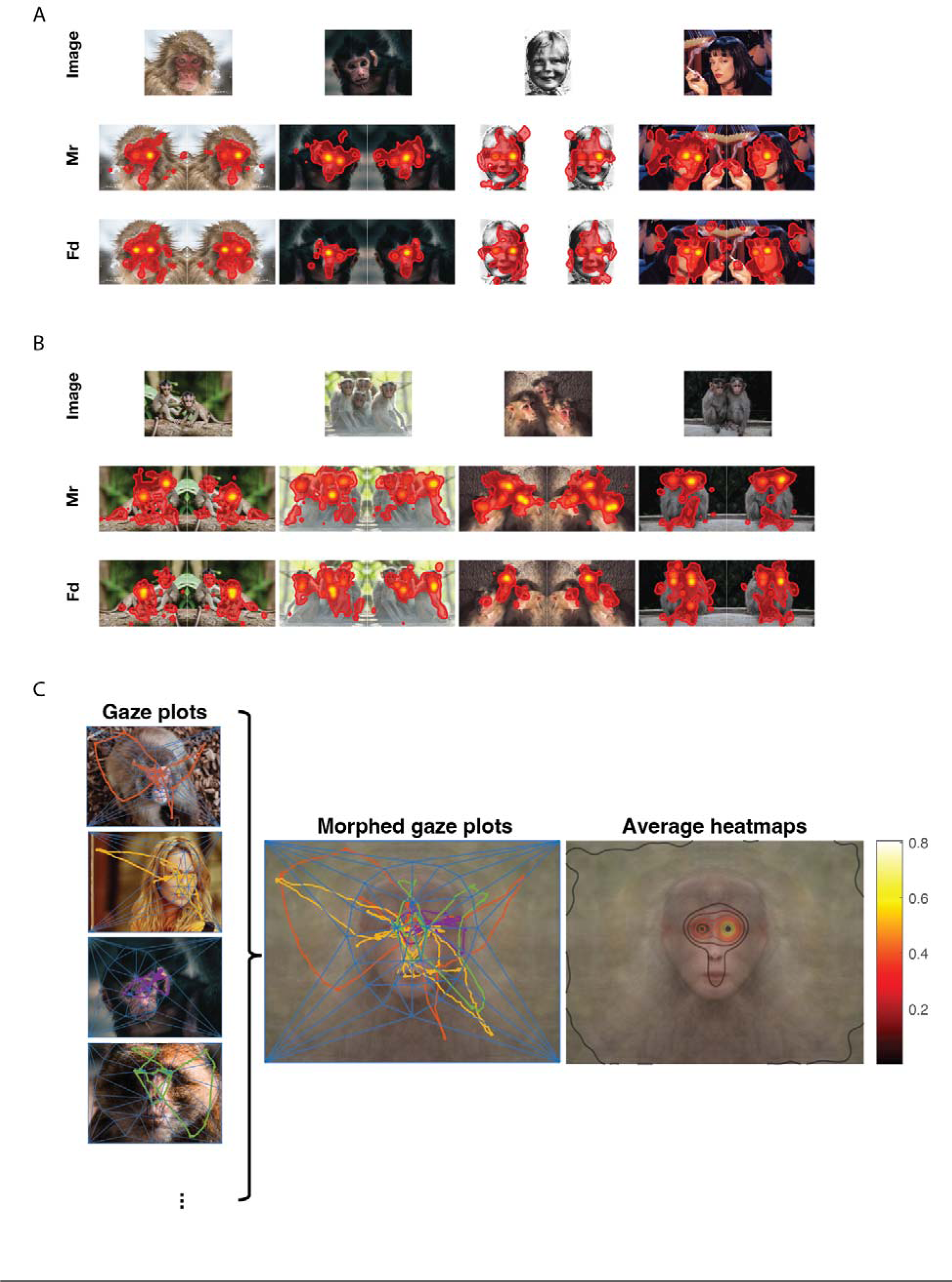
Eye movement patterns for visual stimuli containing faces in baseline sessions. (A-B) Gaze heatmaps were generated to illustrate the typical eye movement patterns of monkeys Mr and Fd during baseline trials for images with a single face (A) and multiple faces (B). Heatmaps represent fixation gaze positions for each image revealing that the monkeys primarily directed their gaze towards the faces and exhibited a strong focus on the eyes for both human and nonhuman primate faces. (C) Morphing was used to average gaze positions and heatmaps across multiple trials. The left column depicts gaze positions for a few exemplar trials in which a face was presented. The blue lines indicate the triangulation on corresponding points for each face. The middle panel illustrates an average face created by warping and averaging all the faces observed by the monkeys, along with the corresponding gaze positions morphed using the same affine transformation matrix. The right panel displays the average gaze heatmaps for all baseline trials presented to monkey Mr, demonstrating the Yarbus-T effect. The results showed a bias towards fixating on the right eye. The human face photographs were obtained from Eye Movements and Vision (17), Pulp Fiction (IMDb link: https://www.imdb.com/title/tt0110912/), and Kill Bill: Vol. 1 (IMDb link: https://www.imdb.com/title/tt0266697/).

Does the fixation pattern follow the Yarbus T? Systematic investigation of gaze patterns offers a technical challenge; how to compare patterns of eye movement across multiple faces? To study this within the entire category of primate faces (including human, ape and monkey faces), we employed face morphing techniques that utilize corresponding points to generate an average gaze heatmap, by merging the data from all trials. Firstly, we identified the corresponding points on each face to construct an average primate face shape. We then used triangulation to determine the affine transform matrix, which was applied to warp gaze positions (see Methods). An average gaze heatmap was computed across all trials in which a face was displayed (Figure 2C). Analysis of the average gaze heatmap from baseline trials revealed that the monkeys primarily directed their gaze towards the interior of the presented faces, with particular emphasis on the eyes, nose, and mouth. However, both monkeys exhibited a noticeable preference for the right side of the faces. We corrected this bias for eye movement patterns within faces during injection trials by subtracting the average gaze heatmap obtained from baseline trials. These results are consistent with earlier findings in macaque monkeys (22).

### The effect of neural inactivation on free viewing of faces

After injecting 10 µl of muscimol in the middle face patch, the monkeys developed a strong preference for looking at the ipsilateral eye compared to the one contralateral to the injection hemisphere. Figure 3A demonstrates changes in the gaze heatmap for a typical image for monkey Mr, highlighting a greater fixation on the ipsilateral side of the face relative to the contralateral side. This trend is also evident in the average gaze heatmaps gathered from all trials featuring primate faces on the screen (Figure 3). To statistically analyze this finding, we defined a rectangular area around each eye on the average face and calculated the time that the animals looked at each eye. We then compared the difference in time spent fixating on each eye. The results showed that after injection of 10 µl of muscimol in the middle face patch, the monkeys on average spent 1.4 and 1.0 s more on the ipsilateral eye (permutation test with 10,000 resampled; *p* < 0.001 and < 0.001; respectively for monkey Mr and Fd). Injection of 10 µl of muscimol in the control area created a similar but significantly weaker effect for monkey Fd (permutation test with 10,000 resampled; *p* = 0.028) and no effect for Monkey Mr (0.1 and 0.4 s difference; permutation test with 10,000 resampled; *p* = 0.389 and 0.003 respectively for monkeys Mr and Fd). Analysis of 5 µl of muscimol did not show any significant level of change in fixation (permutation test with 10,000 resampled; *p* = 0.070 to 0.908 for all comparisons).

**Figure 3:**
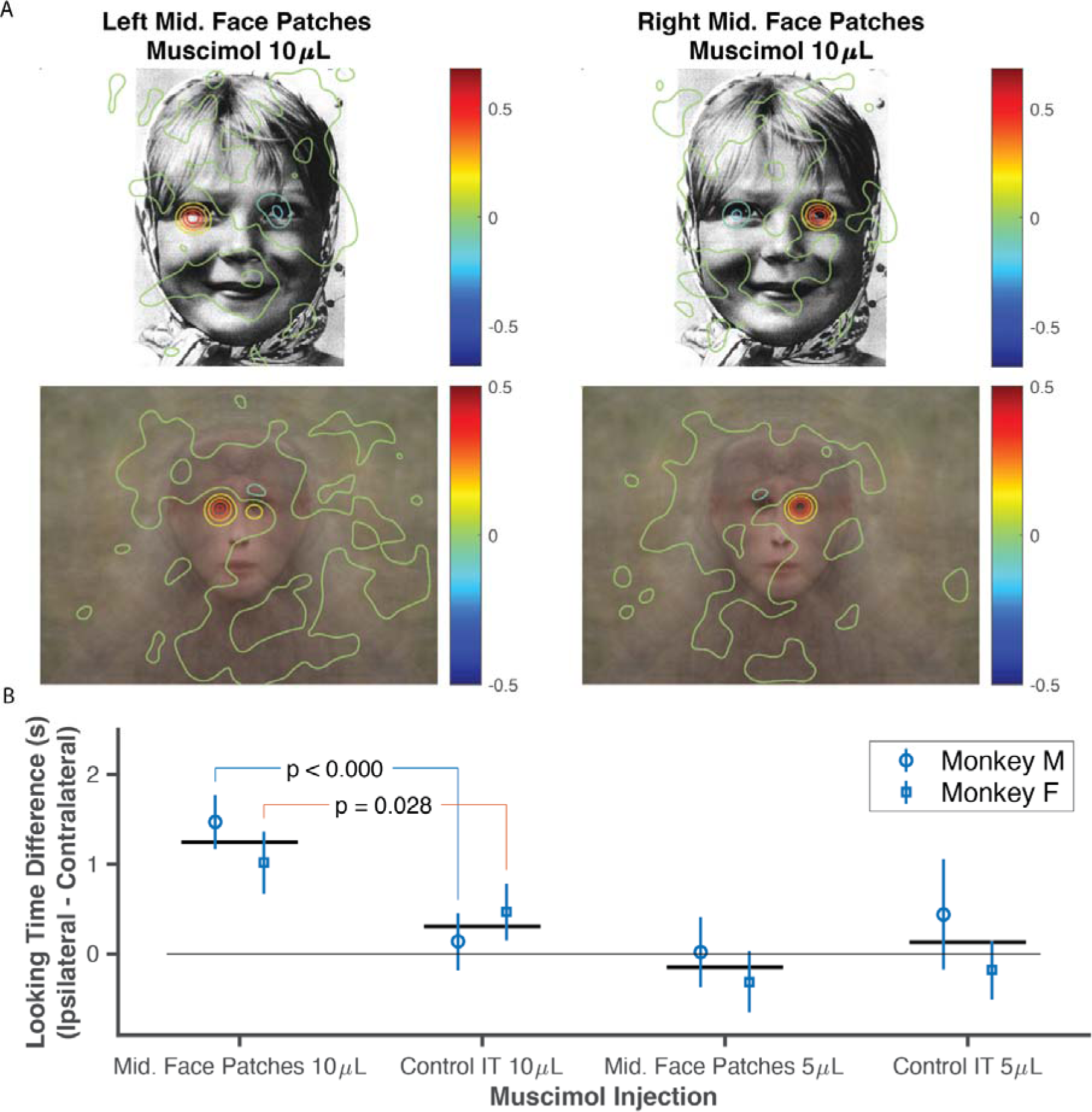
The effect of unilateral neural inactivation in IT cortex on eye movement traces during viewing of faces. The animals exhibited a relative increased duration of gaze towards the eye ipsilateral to the muscimol injection side. (A) The average change in fixation pattern for a typical face image (top) and average of morphed faces (bottom) after 10 µL muscimol injections in the left and right middle face patches for monkey Mr. The heatmaps indicate a greater focus on the ipsilateral eye (with respect to the injection side) compared to the contralateral one. (B) The difference in looking time for the ipsilateral and contralateral eye under different injection conditions for each animal. The error bars represent the 95% confidence intervals, and the horizontal solid lines indicate the average across the animals. The human face photograph was obtained from Eye Movements and Vision (17).

### The effects on free viewing non face objects

Analyzing the potential effects of IT inactivation on looking patterns for nonface objects presents a technical challenge. It is not possible to define corresponding points on nonface objects thus, unlike faces, we cannot morph them into an average form. Therefore, to investigate the potential effect of IT inactivation on nonface objects, we analyzed eye movement on each image separately. This takes a toll on the statistical power of the analysis as we cannot pool the results for nonface objects. Moreover, the existence of Yarbus T for faces allows performing hypothesis testing in search of a potential change of eye movement pattern as shown earlier. However, this is not the case for nonface objects, which leaves only the possibility of searching for a general change in gaze patterns. Given these two limitations, it is not possible to firmly reject a possible effect on looking patterns for nonface objects, even though we did not find any evidence for it.

To conduct statistical analysis on changes in gaze heatmaps after muscimol injections, we computed a pattern change index. This index is the normalized variance of euclidean distance between the gaze patterns of baseline and post-injection trials for each image (see Methods section for more details). Figure 4A displays the average pattern change index for each condition. Our results indicate that the pattern change index was the highest for faces following injection, and we observed a significant difference between the face images when muscimol was injected into the middle face patches (permutation test with 10,000 resamples, p < 0.001 for all comparisons). However, the largest observed effect size was for faces in the case of muscimol injection in the face patches (permutation test with 10,000 resamples, p < 0.001 for all comparisons).

**Figure 4:**
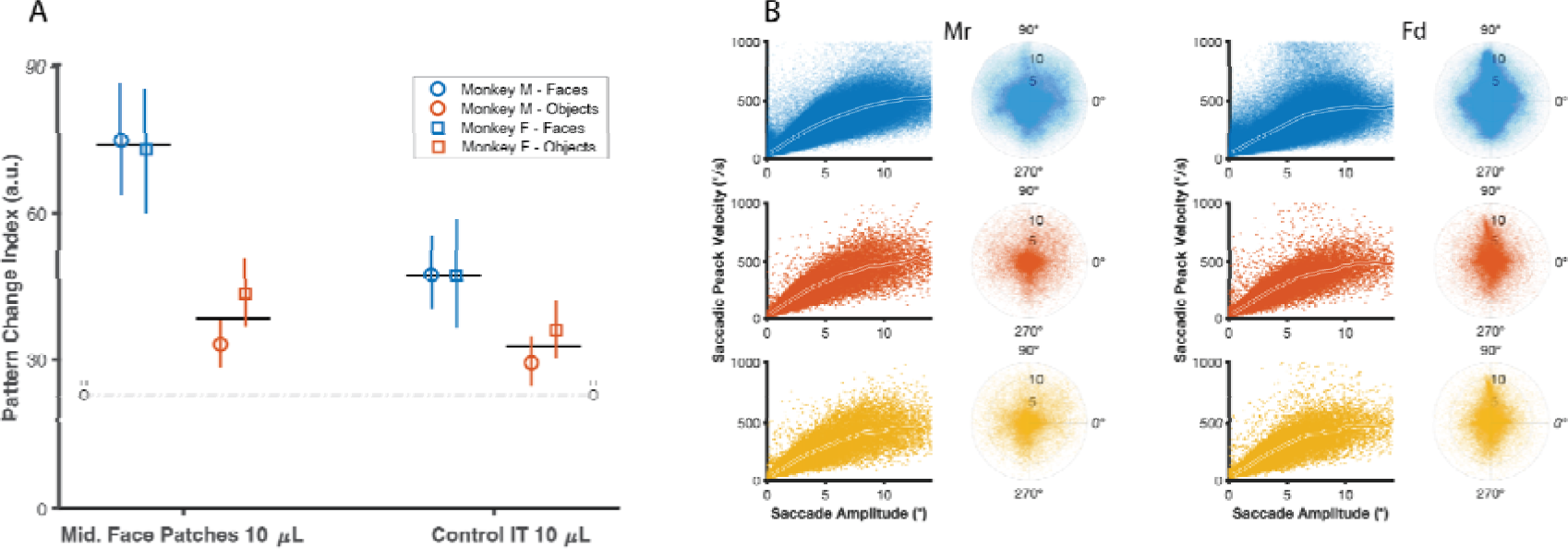
Pattern change index and saccadic properties before and after muscimol injections. (A) This plot illustrates pattern change index across injection conditions. The error bars indicate the 95% confidence intervals, and the solid horizontal lines represent the average across the animals. The dashed bars represent the expected natural variation in the patterns of eye movements estimated from baseline sessions. (B) Saccade properties remained unaffected following muscimol injection. Main sequence and saccade landing points for baseline conditions (blue), as well as 10 µL muscimol injections in the left (red) and right (yellow) middle face patches for both monkeys in trials with a face image. No significant differences were observed, between the main sequences and saccade landing points after muscimol injection.

### The effects on motor features of saccades

To assess the impact of muscimol injections on motor features of saccadic behavior, we examined certain general saccade properties acquired from baseline and 10 µL middle face patch trials. Figure 4B shows the main sequence and polar plot of saccadic landing points for both baseline and sessions conducted after the administration of muscimol in the middle face patch. A regression analysis (see Methods) showed that the saccadic peak velocity followed the main sequence patterns after muscimol injections in the face patches (*R^2^* = 0.72, 0.68 for monkey Mr and 0.69, 0.69 for monkey Fd, respectively for right and left hemispheres). This finding indicates that the oculomotor system retains its capacity to plan and execute saccades effectively.

## Discussion

In line with Yarbus predictions, the present results revealed that faces are behaviorally salient stimuli for the visual system. Morphing the gaze patterns on multiple faces allows us to confirm that free viewing gaze patterns of monkeys naturally follow the Yarbus T. Furthermore, inactivation of face selective neurons in IT cortex alters the patterns of eye movements during free viewing of faces. This intervention does not affect the ability to saccade to faces in the periphery. However, when looking directly at faces, inactivating face selective neurons causes the animals to spend more time viewing the eye ipsilateral to the inactivated side; trimming the contralateral branch of the Yarbus T. These findings also indicate that inactivation of similar volumes of IT cortex outside the face selective area, and inactivation of smaller volumes of cortex (inside or outside the face area) does not induce a comparable effect. We did not observe any specific changes in viewing patterns over nonface objects, although we cannot reject this possibility and we will further discuss it later in this section. These results indicate that the activity of IT face selective neurons is a part of the causal chain for eye movement control. Nevertheless, this causal role is not a direct motor command; the effects depend on the existence of a face in the visual field, also, they do not alter low level motor properties of saccades. Instead, the causal role of face selective neurons is rather shaping the saliency profile that steers the gaze.

Are the observed effects consistent with the physiological response properties of IT neurons? Traditionally, it is assumed that IT neurons, including face selective units, have large bilateral receptive fields even as wide as ∼30 degrees (11, 34). If so, it would be difficult to explain the lateralization and topographical features of the observed results. However, multiple lines of evidence challenge the assumption of large bilateral receptive fields for IT units. Firstly, the size of IT receptive field depends on the way it is measured (35) and more recent studies report sizes as small as 2.5 degrees of visual angle (36, 37). Nevertheless, there is general agreement that typical IT receptive fields are foveally biased and most IT neurons do not respond to far peripheral stimuli. Secondly, IT responses are not bilaterally symmetric and most of the drive comes from the contralateral visual field (36). More importantly, the causal contribution of IT face selective cells is limited to the stimuli presented in the contralateral hemifield (31), at least for the case of a face discrimination task.

Given the topographical properties of IT receptive fields, we can estimate the activity of face-selective neurons when fixating on an eye of a face. Figure 5A presents this estimation when the monkey fixates on the right eye (with respect to the observer) of a face. In this gaze arrangement, the foveated eye (right) is present in the receptive field of face neurons of both hemispheres but the other eye (left) and most of the facial features appear only in the receptive field of the right hemisphere face cells. Therefore, it is reasonable to assume that face neurons of the right hemisphere are more active when looking at the right eye and vice versa. If IT activity serves as a high-level saliency map for the eye movements, the gaze is expected to be directed towards the receptive field of the more active cells which is the left hemifield. This is similar to how saliency functions in low-level visual areas, when the gaze is attracted to the receptive field of the cells that are driven by-say-a flash of light. Now, when face cells are inactivated in the right hemisphere, the balance reverses leading to more fixation time on the right eye according to this hypothetical framework. Here we present this framework only to help organize the current findings, but direct evaluation of this framework requires follow up studies including systematic recording of the neural activity in the absence of inactivation as well as microstimulation of face selective neurons during free viewing of faces.

**Figure 5:**
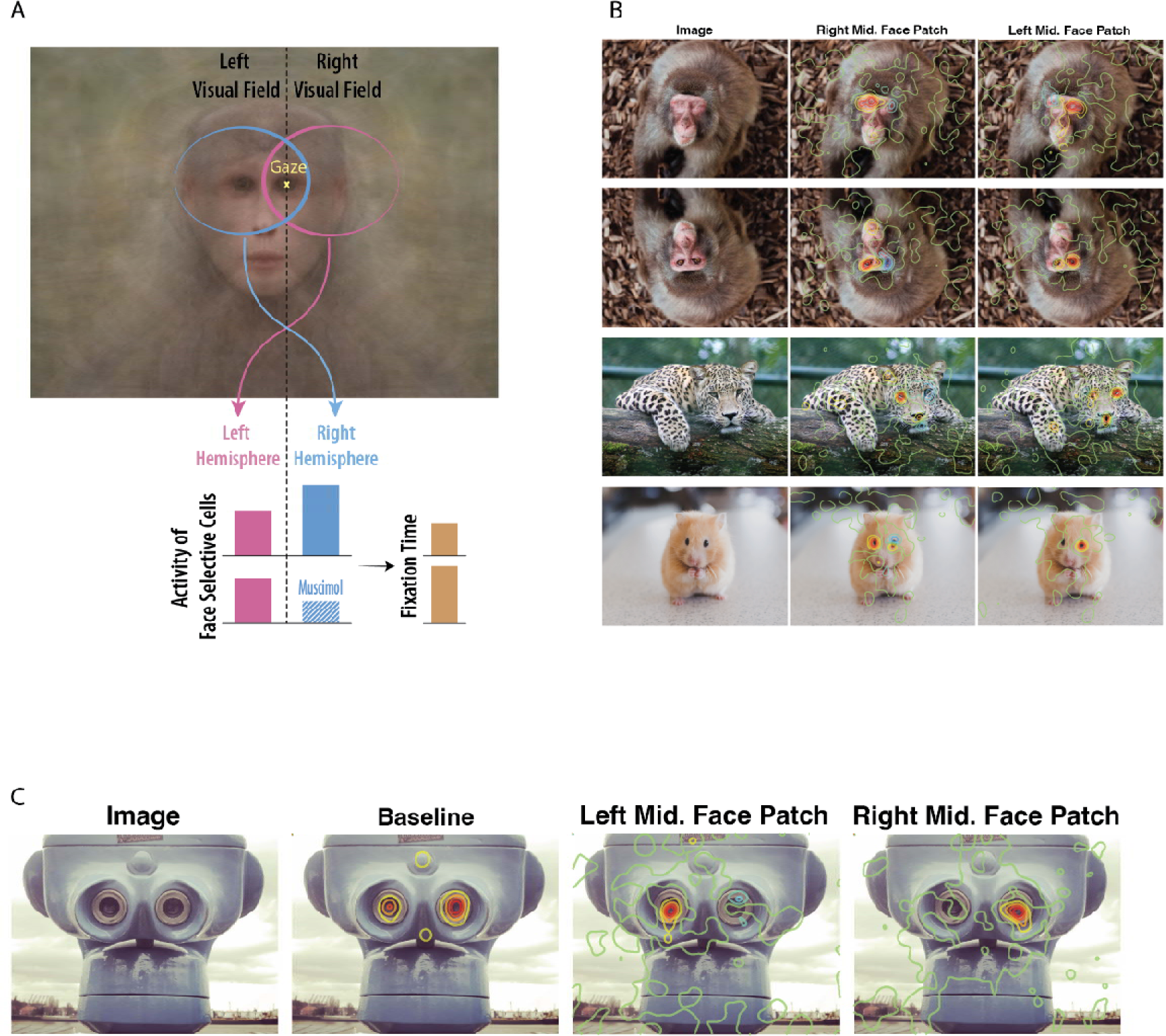
A schematic framework for understanding the effects and a few examples of boundary conditions. (A) We do not have neural recording data available in this study, this framework is merely a schematic, provided based on what is known about neural receptive fields in IT cortex to put the observed effects in neural context. Given the foveal and contralateral biases of IT neurons, it is expected that during fixation on the right eye (represented by the yellow “x”), the receptive fields of face-selective neurons in the right hemisphere encompass both eyes. In contrast, the left hemisphere’s receptive fields include only one eye, less facial features, and a smaller part of the face. Consequently, this configuration is expected to result in increased activity levels in the face-selective neurons of the right hemisphere. Muscimol injection would suppress these activities and invert the balance, ultimately leading to a prolonged fixation time. (B) Average changes in eye movement patterns are provided for examples of boundary condition images for monkey Mr. (C) Average baseline and change in eye movement patterns are presented for face pareidolia for monkey Mr.

How does inactivation of face neurons alter the gaze patterns for direct viewing but not for finding the faces in the periphery? We did not inactivate all of the face neurons even within a hemisphere and it is possible that the activity of the remaining units is sufficient for the search for faces in the periphery. Alternatively, it is possible that the mechanisms underlying finding faces in the periphery and viewing the features within faces are separate from each other. According to this possibility low level saliency of faces presented in the periphery is enough to attract the gaze. Experimental rejection of the first scenario requires inactivation of all face neurons. Nevertheless, performing multiple and prolonged injections offers a technical challenge due to safety considerations and this experiment has to wait for more nimble techniques of neural inactivation. One possible candidate for this purpose is the use of DREADDs (38, 39) where virus injections can be performed in separate sessions but they can all be activated simultaneously.

To explore the boundary conditions, we added a few altered images to the imageset including upside down faces and faces without eyes or with the eyes placed out of the facial frame. Even though this was not the central focus of the current study, we decided to report some of these anecdotal observations as they may inform future experiments. Figure 5B illustrates the effect of inactivation of face selective units on some of the altered images after injection of 10 uL of muscimol in the face patch. In the case of inverted face condition, we were curious to see whether there is a similar lateralized effect and if so, which side of the face would be ignored. An object-centered representation of faces (40, 41) predicts flipping the effect side, in contrast, a head-centered representation (42–44) anticipates persistence of the effect on the same side with respect to the observer. The results were consistent with a head-centered basis for cortical representation of faces. The other image alteration tested here was to eliminate the eyes on a monkey face and cover their place with face skin. This alteration was inspired by two facts. First, our effects seem to be mainly focused on the eyes of a viewed face. Second, previous studies suggest that the eyes are the main driver of the majority of face selective units in the posterior face patch (45) as well as anterior medial face patch (46). Looking patterns for these unusually altered faces elicited comparable effects to unaltered faces; the animals spent more time looking at where the ipsilateral eye was supposed to be and ignored the corresponding contralateral position. This shows that an “eye” is not defined for the face processing system merely by its internal features such as a round pupil and that contextual cues can also inform the system (47).

Another boundary condition tested here was the case of face pareidolia. The phenomenon of perceiving faces in objects, known as face pareidolia, is not limited to humans; it also happens to rhesus monkeys (48). Figure 5C, the bottom row, depicts a case of such a phenomenon, a tower-viewer that resembles a face. Consistent with Taubert et al. (48) the monkeys’ eye movement patterns resembled a Yarbus T where the binoculars of the tower-viewer corresponded to the eyes of a face. Is this pareidolia effect mediated through the face selective neurons? While correlational involvement of face selective parts of the cortex in face pareidolia has been shown (49), our results for the first time establish a causal link between IT face selective units and face pareidolia.

Is IT activity generally used to drive eye movements or is it a special case for faces? Our findings did not indicate any systematic change in eye movement patterns for objects following muscimol injection in IT cortex. However, we cannot reject the possibility of such changes entirely. Unlike faces, objects lack corresponding points, making it impractical to systematically aggregate gaze heatmaps across multiple objects. Consequently, the statistical power to detect a significant effect in the case of objects is diminished. Moreover, given the clustering of face selective neurons in the cortex, muscimol would likely affect neurons with similar properties, augmenting the behavioral impact of inactivation. As a result, the causal role of other IT neurons in eye movements for nonface objects remains a possibility for further exploration using more specific techniques. For instance, molecular labeling and perturbation of object selective neuronal populations that are not spatially clustered (50–55) can follow up this question.

In conclusion, the findings presented in this study establish a causal relationship between the activity of face-selective neurons in IT cortex and the oculomotor system. These results indicate how high-level visual processing can determine the future input to the visual system by steering the eyes towards the relevant parts of the visual field. They also highlight the potential utility of eye movements as a valuable tool for investigating visual processing within high-level visual areas.

## Methods

In this study, we collected free viewing eye movement data from two adult male rhesus monkeys (Macaca mulatta), designated as Mr and Fd. All procedures were carried out in accordance with the guidelines of the National Institute of Mental Health Animal Use and Care Committee.

### Functional localization Data acquisition

To identify the discrete regions or patches of inferior temporal cortex that responded preferentially to faces, we used an independent functional localizer experiment (56–60). While the subjects were awake and fixating, we presented images (30 greyscale images per category) of 6 different categories of visual stimuli (human faces, monkey faces, scenes, objects, phase scrambled human faces and phase scrambled monkey faces). The order of blocks, as well as image order within each block, were randomized and presented using Psychtoolbox (61, 62) and PLDAPS toolbox (63) on MATLAB (MathWorks, version R2018b). Stimuli were cropped images presented on a square canvas that was 12 dva in height. All six categories were presented in each run in a standard on/off block design (12 blocks in total). Each block lasted for 16.5 s. During a “stimulus on” block, 15 images were presented one at a time for 900 ms and were followed by a 200 ms inter-stimulus interval (ISI). Eye position throughout the scan sessions was recorded using a magnetoresistance-compatible infrared camera (MRC Systems, Heidelberg, Germany) at 60 Hz. For maintaining fixation within a central fixation window with a 4 deg diameter, subjects received juice rewards (average juice intervals varied between 0.6 −3 s). We removed any run from the analysis where the monkey did not fixate within the fixation window for more than 70% of the time.

Functional data were acquired in a 4.7T Bruker Biospin scanner with a vertical bore (Bruker Biospec, Ettlingen, Germany). Before each scanning session, a contrast agent (monocrystalline iron oxide nanocolloid; MION) was injected into the femoral vein to increase the hemodynamic response. MION doses were determined independently for each subject (∼8 to 10 mg/kg). We collected whole-brain images with a four-channel transmit-and-receive radiofrequency coil system (Rapid MR International, Columbus, OH). A low-resolution anatomical scan was also acquired in the same session to serve as an anatomical reference [modified driven equilibrium Fourier transform (MDEFT) sequence: voxel size, 1.5 mm by 0.5 mm by 0.5 mm; field of view (FOV), 96 mm by 48 mm; matrix size, 192 × 96; echo time (TE), 3.95 ms; and repetition time (TR), 11.25 ms]. Functional echo planar imaging (EPI) scans were collected as 42 sagittal slices with an in-plane resolution of 1.5 mm by 1.5 mm and a slice thickness of 1.5 mm. The TR was 2.2 s, and the TE was 16 ms (FOV = 96 mm x 54 mm; matrix size, 64 × 36 m; flip angle, 75°). For anatomical registration and MR targeting, we also acquired high-resolution T1-weighted whole-brain anatomical scans under sedation in a 4.7T Bruker scanner with an MDEFT sequence. Imaging parameters were as follows: voxel size, 0.5 mm by 0.5 mm by 0.5 mm; TE, 4.9 ms; TR, 13.6 ms; and flip angle, 14°. These scans were used to create a high-resolution template for each subject.

### fMRI preprocessing and localization

All EPI data were analyzed using AFNI software (http://afni.nimh.nih.gov/afni) (64). Raw images were first converted from Bruker into AFNI data file format. The data collected in a single session were corrected for static magnetic field inhomogeneities using the PLACE algorithm (65). The time series data were then slice time– corrected and realigned to the volume with the minimum outliers. All the data for a given subject were aligned to the corresponding high-resolution template for that subject, allowing for the combination of data across multiple sessions. The first two volumes of data in each EPI sequence were discarded. The volume-registered data were then spatially smoothed with a 3-mm Gaussian kernel and rescaled to reflect percentage signal change from baseline. The statistical significance values of the differential signal changes from the functional scans were projected onto the high-resolution anatomical scan and surface model for visualization. The face-selective patches in ITC were identified in all subjects using the following contrast: activations evoked by (human faces + monkey faces) > activations evoked by (scenes + objects; see Figure 1A-D). Targeting of the mediolateral face patches were based on the corresponding peak activations in the middle STS region. The targeted control areas were chosen on the convexity of the IT cortex 4 to 5.5 mm anterior and 4 to 8 mm ventral to the middle face patches where no face selectivity was observed.

### Free viewing experiment Apparatus

The experiments were conducted in a dark room using NIMH MonkeyLogic (66) on MATLAB (MathWorks, version R2019a). The animals were positioned 57 cm away from a 32-inch, 1920×1080 pixel, 120 Hz, LCD Monitor (Display++, Cambridge Research System). Eye tracking was performed using monocular corneal reflection and pupil tracking (Eyelink 1000 Plus, SR Research) at a frequency of 1000 Hz.

### Behavioral Experiments

Each session began with a calibration procedure consisting of 13 points at the start of each session. The monkeys initiated a trial by fixating for 1 s on a central fixation point (a black circle with a radius of 0.3°) against a gray background. An image (28° × 21°) was then displayed on the screen for 10 s. The monkeys received liquid reward every 1 to 3.5 s as long as they maintained their gaze on the image. Any breaks in looking at the image that lasted less than 500 ms were ignored, allowing the animals to blink naturally.

Only the trails that the animals spent more than 7 s were included for further analysis. The image was randomly selected from a set of 62 unique images, along with their left-right flipped versions, making a total of 128 images. The animals performed 129 and 78 baseline sessions, and 28 and 23 muscimol injection sessions respectively for monkey Mr and Fd. Table xyz illustrates the number of injections on each site and volume for each monkey. On average, the animals performed 135.7 and 321.1 trials in each session respectively for monkey Mr and Fd.

**Table 1:**
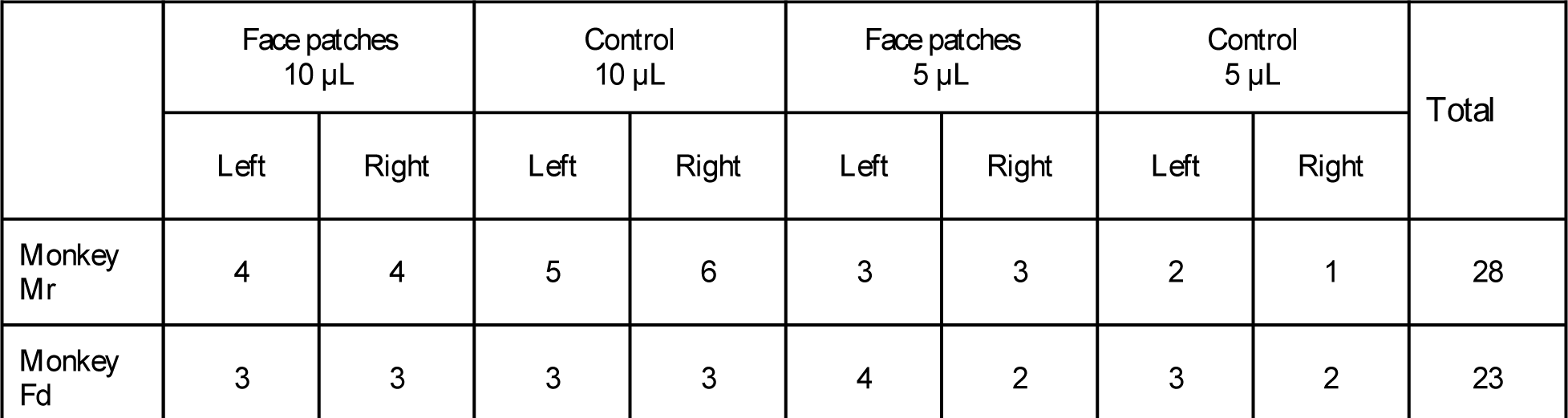
Number of injection sessions for each condition in each animal

### Injection apparatus and parameters

For microinjections, a handmade injection circuit was used, based on the design presented by Noudoost and Moore (67); a 30-gauge (0.304 μm outer diameter) stainless steel lancet point cannula with a long 15° primary bevel, coupled with a 5-25 µL gas tight syringe (1700 series, Hamilton), was used for drug delivery. For infusion, we used a syringe pump (Pump 11 Elite, Harvard Apparatus). For cannula insertion and extraction, a 3d-printed grid and an oil hydraulic micromanipulator (MO-952, Narishige) was used for accurate targeting and minimal damage to the cortex. To prepare the muscimol solution, a commonly used GABA_A_ agonist, muscimol powder (Sigma-Aldrich) was diluted in normal saline to a concentration of 5 mg/mL and then filtered through a sterile filter.

Injections were performed in both the left and right hemispheres through a rectangular cranial chamber measuring 45 × 25 mm (coronal × sagittal dimensions). The chamber was centered at the sagittal midline and approximately 8 mm anterior-posterior stereotaxic coordinate. During the high-resolution T1-weighted whole-brain anatomical scan, the cranial chambers were filled with a clinical-grade MR contrast agent Gadolinium-DTPA, which was diluted with normal saline to 5 mmol/L. This enabled us to identify the four corners of the grid positioned within the chamber. We then selected the appropriate grid hole and depth for each target position based on the angle of grid holes 30° by fitting a plane on the four corners.

To confirm the accuracy of the targeting procedure, we infused 1 µL Gadolinium-DTPA diluted with normal saline to 5 mmol/L. The cortical spread of Gadolinium-DTPA is similar to that of muscimol, as shown in previous studies (68), and can be detected in an anatomical MRI scan. We assessed the overlap of Gadolinium-DTPA spread with the functionally defined peak activations by aligning the post-injection anatomical volume and functional localizer data in a post-hoc analysis (see Figure 1B-D).

### Saccade detection

To detect saccades, we applied a velocity threshold criterion of six standard deviations from the median saccade velocity during fixation at the beginning of each trial. The start and end points of saccades were determined by marking the points where velocity fell below two standard deviations from the median saccade velocity during fixation.

### Fixation heatmaps

To generate the fixation heatmaps, we excluded saccades greater than 1 dva from the eye movement traces. Next, we superimposed all the eye movement traces from each trial onto a heatmap grid. We then applied a 2d Gaussian filter to the heatmaps, with a standard deviation of 0.5 dva.

### Morphing

To average facial features and analyze eye movement patterns on the primate faces presented during the experiment, we start by manually identifying 26 corresponding points on each face. The average face shape is then determined by calculating the mean *x* and *y* coordinates for each corresponding point. To create corresponding triangles on each face, we use the Delaunay Triangulation.

For each triangle, we can create an affine transform using the following equation:

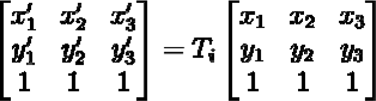

where ***x***s and ***y***s represent the vertices of a triangle on the original image and ***x’***s and ***y’***s represent the vertices for the corresponding triangle on the morphed image. ***T*i** is the affine transform matrix for triangle ***i*** that can be calculated using the above formula. This allows us to transform any point **(*x,y*)** from the original image within the triangle ***i***, to new coordinates on the new image using the following equation:

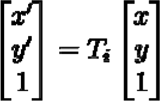

### Variance of gaze heatmaps after injection

To check the effect of injection, we calculated the variance of gaze heatmaps before and after injection. For this we resampled heatmaps to 140×105 pixels (providing 5 pixels per degree in visual angle), then applied a 2d gaussian convolution with standard deviation of 2.5 pixels. Then for each trial we defined a pattern change index:

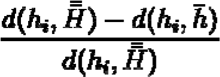

Where ***d*** is the euclidean distance between two heatmaps. ***h_i_*** is gaze heatmap for trial 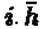 is the average gaze heatmap for the visual stimulus across all trials in injection condition. 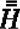 is the grand average of gaze heatmaps across all the trials in the injection condition and all the baseline trials for the same visual stimulus. The euclidean distance between two heatmaps (e.g. ***u*** and ***v***) can be calculated by the following equation:

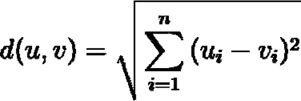

### Analysis of saccadic main sequence

We employed regression analysis to fit a polynomial function to saccadic properties derived from trials in baseline sessions and each injection condition, treating each animal separately. The polynomial function used was as follows:

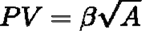

Here ***PV*** represents the saccadic peak velocity, ***A*** denotes the saccadic amplitude, and β is the regression coefficient. The regression analysis yielded substantial results, with R-squared values ranging from 0.65 to 0.72 for monkey Mr and 0.56 to 0.69 for monkey Fd.

## Acknowledgements

We thank David Yu for his crucial contributions to the surgeries and MRI scanning. This research was supported by the Intramural Research Program of the NIMH ZIAMH002958 (to A.A.).

## References

1. L. Itti, C. Koch, E. Niebur, A model of saliency-based visual attention for rapid scene analysis. IEEE Trans. Pattern Anal. Mach. Intell. 20, 1254–1259 (1998).

2. L. Itti, C. Koch, A saliency-based search mechanism for overt and covert shifts of visual attention. Vision Res. 40, 1489–1506 (2000).

3. X. Hou, L. Zhang, Saliency detection: A spectral residual approach in 2007 *IEEE Conference on Computer Vision and Pattern Recognition*, (IEEE, 2007) https://doi.org/10.1109/cvpr.2007.383267.

4. A. Treisman, S. Gormican, Feature analysis in early vision: evidence from search asymmetries. Psychol. Rev. 95, 15–48 (1988).

5. M. Zhang, et al., Finding any Waldo with zero-shot invariant and efficient visual search. Nat. Commun. 9, 3730 (2018).

6. M. Zhang, et al., Look twice: A generalist computational model predicts return fixations across tasks and species. PLoS Comput. Biol. 18, e1010654 (2022).

7. M. Kümmerer, M. Bethge, T. S. A. Wallis, DeepGaze III: Modeling free-viewing human scanpaths with deep learning. J. Vis. 22, 7 (2022).

8. A. Borji, L. Itti, State-of-the-art in visual attention modeling. IEEE Trans. Pattern Anal. Mach. Intell. 35, 185–207 (2013).

9. M. Kümmerer, L. Theis, M. Bethge, Deep Gaze I: Boosting Saliency Prediction with Feature Maps Trained on ImageNet (2014) (April 9, 2023).

10. E. L. Schwartz, R. Desimone, T. D. Albright, C. G. Gross, Shape recognition and inferior temporal neurons. Proc. Natl. Acad. Sci. U. S. A. 80, 5776–5778 (1983).

11. R. Desimone, T. D. Albright, C. G. Gross, C. Bruce, Stimulus-selective properties of inferior temporal neurons in the macaque. J. Neurosci. 4, 2051–2062 (1984).

12. K. Tanaka, H. Saito, Y. Fukada, M. Moriya, Coding visual images of objects in the inferotemporal cortex of the macaque monkey. J. Neurophysiol. 66, 170–189 (1991).

13. D. Y. Tsao, M. S. Livingstone, Mechanisms of face perception. Annu. Rev. Neurosci. 31, 411–437 (2008).

14. D. I. Perrett, E. T. Rolls, W. Caan, Visual neurones responsive to faces in the monkey temporal cortex. Exp. Brain Res. 47, 329–342 (1982).

15. A. Afraz, R. Kiani, H. Esteky, Microstimulation of inferotemporal cortex influences face categorization. Nature 442, 692–695 (2006).

16. D. A. Leopold, A. J. O’Toole, T. Vetter, V. Blanz, Prototype-referenced shape encoding revealed by high-level aftereffects. Nat. Neurosci. 4, 89–94 (2001).

17. A. L. Yarbus, Eye Movements During Perception of Complex Objects. Eye Movements and Vision, 171–211 (1967).

18. S. W. Janik, A. R. Wellens, M. L. Goldberg, L. F. Dell’Osso, Eyes as the center of focus in the visual examination of human faces. Percept. Mot. Skills 47, 857–858 (1978).

19. J. J. S. Barton, N. Radcliffe, M. V. Cherkasova, J. Edelman, J. M. Intriligator, Information processing during face recognition: the effects of familiarity, inversion, and morphing on scanning fixations. Perception 35, 1089–1105 (2006).

20. J. Arizpe, V. Walsh, G. Yovel, C. I. Baker, The categories, frequencies, and stability of idiosyncratic eye-movement patterns to faces. Vision Res. 141, 191–203 (2017).

21. R. R. Althoff, N. J. Cohen, Eye-movement-based memory effect: a reprocessing effect in face perception. J. Exp. Psychol. Learn. Mem. Cogn. 25, 997–1010 (1999).

22. K. Guo, R. G. Robertson, S. Mahmoodi, Y. Tadmor, M. P. Young, How do monkeys view faces?-A study of eye movements. Exp. Brain Res. 150, 363–374 (2003).

23. I. N. Springer, et al., [Facial aesthetics part I-the significance of the triangle of yarbus]. Mund Kiefer Gesichtschir. 11, 145–151 (2007).

24. R. Cappelli, A. Franco, D. Maio, Gabor Saliency Map for Face Recognition in *14th International Conference on Image Analysis and Processing (ICIAP 2007)*, (IEEE, 2007) https://doi.org/10.1109/iciap.2007.4362818.

25. G. A. R. Johnston, Muscimol as an ionotropic GABA receptor agonist. Neurochem. Res. 39, 1942–1947 (2014).

26. J. K. Hesse, D. Y. Tsao, The macaque face patch system: a turtle’s underbelly for the brain. Nat. Rev. Neurosci. 21, 695–716 (2020).

27. D. Y. Tsao, W. A. Freiwald, T. A. Knutsen, J. B. Mandeville, R. B. H. Tootell, Faces and objects in macaque cerebral cortex. Nat. Neurosci. 6, 989–995 (2003).

28. N. Kanwisher, G. Yovel, The fusiform face area: a cortical region specialized for the perception of faces. Philos. Trans. R. Soc. Lond. B Biol. Sci. 361, 2109–2128 (2006).

29. N. Kanwisher, Domain specificity in face perception. Nat. Neurosci. 3, 759–763 (2000).

30. P. L. Aparicio, E. B. Issa, J. J. DiCarlo, Neurophysiological Organization of the Middle Face Patch in Macaque Inferior Temporal Cortex. J. Neurosci. 36, 12729–12745 (2016).

31. A. Afraz, E. S. Boyden, J. J. DiCarlo, Optogenetic and pharmacological suppression of spatial clusters of face neurons reveal their causal role in face gender discrimination. Proc. Natl. Acad. Sci. U. S. A. 112, 6730–6735 (2015).

32. D. B. T. McMahon, B. E. Russ, H. D. Elnaiem, A. I. Kurnikova, D. A. Leopold, Single-unit activity during natural vision: diversity, consistency, and spatial sensitivity among AF face patch neurons. J. Neurosci. 35, 5537–5548 (2015).

33. B. E. Russ, D. A. Leopold, Functional MRI mapping of dynamic visual features during natural viewing in the macaque. Neuroimage 109, 84–94 (2015).

34. C. G. Gross, C. E. Rocha-Miranda, D. B. Bender, Visual properties of neurons in inferotemporal cortex of the Macaque. J. Neurophysiol. 35, 96–111 (1972).

35. E. T. Rolls, N. C. Aggelopoulos, F. Zheng, The receptive fields of inferior temporal cortex neurons in natural scenes. J. Neurosci. 23, 339–348 (2003).

36. H. Op De Beeck, R. Vogels, Spatial sensitivity of macaque inferior temporal neurons. J. Comp. Neurol. 426, 505–518 (2000).

37. J. J. DiCarlo, J. H. R. Maunsell, Anterior inferotemporal neurons of monkeys engaged in object recognition can be highly sensitive to object retinal position. J. Neurophysiol. 89, 3264–3278 (2003).

38. J. Bonaventura, et al., High-potency ligands for DREADD imaging and activation in rodents and monkeys. Nat. Commun. 10, 4627 (2019).

39. M. A. G. Eldridge, et al., Chemogenetic disconnection of monkey orbitofrontal and rhinal cortex reversibly disrupts reward value. Nat. Neurosci. 19, 37–39 (2016).

40. M. E. Hasselmo, E. T. Rolls, G. C. Baylis, V. Nalwa, Object-centered encoding by face-selective neurons in the cortex in the superior temporal sulcus of the monkey. Exp. Brain Res. 75, 417–429 (1989).

41. C. R. Olson, Brain representation of object-centered space in monkeys and humans. Annu. Rev. Neurosci. 26, 331–354 (2003).

42. A. Afraz, P. Cavanagh, The gender-specific face aftereffect is based in retinotopic not spatiotopic coordinates across several natural image transformations. J. Vis. 9, 10.1–17 (2009).

43. S.-R. Afraz, P. Cavanagh, Retinotopy of the face aftereffect. Vision Res. 48, 42–54 (2008).

44. J. E. Dickinson, H. K. Mighall, R. A. Almeida, J. Bell, D. R. Badcock, Rapidly acquired shape and face aftereffects are retinotopic and local in origin. Vision Res. 65, 1–11 (2012).

45. E. B. Issa, J. J. DiCarlo, Precedence of the eye region in neural processing of faces. J. Neurosci. 32, 16666–16682 (2012).

46. E. N. Waidmann, K. W. Koyano, J. J. Hong, B. E. Russ, D. A. Leopold, Local features drive identity responses in macaque anterior face patches. Nat. Commun. 13, 5592 (2022).

47. M. J. Arcaro, C. Ponce, M. Livingstone, The neurons that mistook a hat for a face. Elife 9 (2020).

48. J. Taubert, S. G. Wardle, M. Flessert, D. A. Leopold, L. G. Ungerleider, Face Pareidolia in the Rhesus Monkey. Curr. Biol. 27, 2505–2509.e2 (2017).

49. G. Akdeniz, S. Toker, I. Atli, Neural mechanisms underlying visual pareidolia processing: An fMRI study. Pak. J. Med. Sci. Q. 34, 1560–1566 (2018).

50. C. J. Guenthner, K. Miyamichi, H. H. Yang, H. C. Heller, L. Luo, Permanent genetic access to transiently active neurons via TRAP: targeted recombination in active populations. Neuron 78, 773–784 (2013).

51. T. Kawashima, et al., Functional labeling of neurons and their projections using the synthetic activity-dependent promoter E-SARE. Nat. Methods 10, 889–895 (2013).

52. X. Liu, et al., Optogenetic stimulation of a hippocampal engram activates fear memory recall. Nature 484, 381–385 (2012).

53. K. Sakurai, et al., Capturing and Manipulating Activated Neuronal Ensembles with CANE Delineates a Hypothalamic Social-Fear Circuit. Neuron 92, 739–753 (2016).

54. A. T. Sørensen, et al., A robust activity marking system for exploring active neuronal ensembles. Elife 5 (2016).

55. J. H. Hyun, et al., Tagging active neurons by soma-targeted Cal-Light. Nat. Commun. 13, 7692 (2022).

56. J. Taubert, et al., Clutter substantially reduces selectivity for peripheral faces in the macaque brain. J. Neurosci. 42, 6739–6750 (2022).

57. J. Taubert, et al., The cortical and subcortical correlates of face pareidolia in the macaque brain. Soc. Cogn. Affect. Neurosci. 17, 965–976 (2022).

58. J. Taubert, et al., Parallel Processing of Facial Expression and Head Orientation in the Macaque Brain. J. Neurosci. 40, 8119–8131 (2020).

59. J. Taubert, et al., A broadly tuned network for affective body language in the macaque brain. Sci Adv 8, eadd6865 (2022).

60. J. Taubert, S. G. Wardle, L. G. Ungerleider, What does a “face cell” want?’. Prog. Neurobiol. 195, 101880 (2020).

61. D. H. Brainard, The Psychophysics Toolbox. Spat. Vis. 10, 433–436 (1997).

62. D. G. Pelli, The VideoToolbox software for visual psychophysics: transforming numbers into movies. Spat. Vis. 10, 437–442 (1997).

63. K. M. Eastman, A. C. Huk, PLDAPS: A Hardware Architecture and Software Toolbox for Neurophysiology Requiring Complex Visual Stimuli and Online Behavioral Control. Front. Neuroinform. 6, 1 (2012).

64. R. W. Cox, AFNI: software for analysis and visualization of functional magnetic resonance neuroimages. Comput. Biomed. Res. 29, 162–173 (1996).

65. Q.-S. Xiang, F. Q. Ye, Correction for geometric distortion and N/2 ghosting in EPI by phase labeling for additional coordinate encoding (PLACE). Magn. Reson. Med. 57, 731–741 (2007).

66. J. A. Hwang, A. R. Mitz, E. A. Murray, NIMH MonkeyLogic: Behavioral control and data acquisition in MATLAB. J. Neurosci. Methods 323, 13–21 (2019).

67. B. Noudoost, T. Moore, A reliable microinjectrode system for use in behaving monkeys. J. Neurosci. Methods 194, 218–223 (2011).

68. J. D. Heiss, S. Walbridge, A. R. Asthagiri, R. R. Lonser, Image-guided convection-enhanced delivery of muscimol to the primate brain. J. Neurosurg. 112, 790–795 (2010).

